# A homing CRISPR mouse resource for barcoding and lineage tracing

**DOI:** 10.1101/280289

**Authors:** Reza Kalhor, Kian Kalhor, Kathleen Leeper, Amanda Graveline, Prashant Mali, George M. Church

## Abstract

Cellular barcoding using nuclease-induced genetic mutations is an effective approach that is emerging for recording biological information, including developmental lineages. We have previously introduced the homing CRISPR system as a promising methodology for generating such barcodes with scalable diversity and without crosstalk. Here, we present a mouse line (MARC1) with multiple genomically-integrated and heritable homing guide RNAs (hgRNAs). We determine the genomic locations of these hgRNAs, their activity profiles during gestation, and the diversity of their mutants. We apply the line for unique barcoding of mouse embryos and differential barcoding of embryonic tissues. We conclude that this mouse line can address the unique challenges associated with in vivo barcoding in mammalian model organisms and is thus an enabling platform for recording and lineage tracing applications in a mammalian model system.

## Introduction

Cellular barcoding, defined as marking individual cells with unique heritable genetic sequences, is an effective strategy for tracking cells (Gerrits et al. 2010; Naik et al. 2014; Church et al. 2014). Whereas previously these barcodes had to be generated in vitro and delivered to cells by viral or other means of transfection (Walsh & Cepko 1992; Ma et al. 2017), recent advances in genome engineering technologies have enabled in vivo barcode generation. For that, a locus is targeted for rearrangement or mutagenesis such that a diverse set of outcomes are generated in different cells (Peikon et al. 2014). These in vivo-generated barcodes drastically expand the scope of cellular barcoding strategies, promising deep and precise lineage tracing from single-cell to whole-organism level (McKenna et al. 2016; Kalhor et al. 2017; Frieda et al. 2017; Church et al. 2014) and recording of cellular signals over time (Perli et al. 2016; Sheth et al. 2017).

In eukaryotes, nuclease-induced double-stranded DNA breaks (Gaj et al. 2013; Mali et al. 2013) followed by nonhomologous end-joining (NHEJ) that creates mutations during repair (Lieber & Wilson 2010) can be used to generate in vivo barcodes. Multiple studies establish proof of this principle in recording and lineage tracing (Woodworth et al. 2017; Schmidt & Platt 2017; Spanjaard & Junker 2017), with demonstrations in cultured cells (Junker et al. 2016; Frieda et al. 2017; Kalhor et al. 2017; Schmidt et al. 2017; Perli et al. 2016), axolotl (Flowers et al. 2017), and zebrafish (McKenna et al. 2016; Raj et al. 2017). However, no demonstrations have yet been carried out in mouse, a model organism more relevant to human development (Barriga et al. 2015). Moreover, while most studies so far rely on synthetic genetic elements to generate barcodes in vivo, no model organism with integrated barcoding elements has been established to facilitate expanding the use of this technology beyond the labs that specialize in it and to enable the reproduction of results between groups.

Compared to other vertebrate model systems, establishing a line for in vivo barcoding is critically important in mouse because its genetic manipulation is more challenging: unlike fish and salamander, mouse does not feature external fertilization, large brood sizes, and a large one-cell-stage embryo that can be obtained and injected with ease and reliability. These challenges make it difficult to inject and track a large number of individual zygotes in mouse, necessitating genetically integrated barcoding elements, i.e., the nuclease and barcoding loci. On the other hand, mammalian development is more challenging to track than that of other vertebrate model systems: unlike fish and salamander, gestation in mouse transpires inside a living animal, takes longer times, and involves complex extraembryonic structures (Rossant & Cross 2002); it also produces more intermediate cell types and leads to more complex organ systems in many cases (Speck et al. 2002; Sharma & Goffinet 2012). Therefore, the elements in a barcoding line should be well-characterized, reliable, and highly effective to allow for informative analysis on a small number of animals. Furthermore, as modifying and upgrading genetically integrated systems in mouse present a large barrier, such elements would ideally be versatile as well as easily expandable and adaptable to new technologies that emerge.

Here, we present a mouse line for barcoding and lineage tracing applications based on the homing CRISPR system. We have previously described homing CRISPR guide RNAs (hgRNAs) (Kalhor et al. 2017), a modified version of canonical guide RNAs (gRNAs) that target and retarget their own locus (Figure 1a) (Perli et al. 2016). We demonstrated that hgRNAs create a substantially larger diversity of mutants than canonical gRNAs (Figure 1b), can act as expressed genetic barcodes, enable lineage tracing in populations of cultured cells, and can be detected using high-throughput or in situ sequencing (Kalhor et al. 2017). We also argued that the barcode diversity generated by hgRNAs can be expanded exponentially by simultaneous deployment of multiple independently-acting hgRNAs, making them a suitable candidate for addressing the challenges associated with in vivo barcoding in mouse. The mouse line presented here carries multiple genomically integrated hgRNAs which can be activated upon introduction of *S. pyogenes* Cas9. We have extensively characterized these hgRNAs and profiled their behavior during gestation, thereby establishing the mouse line’s suitability for recording and lineage tracing applications.

**Figure 1.**
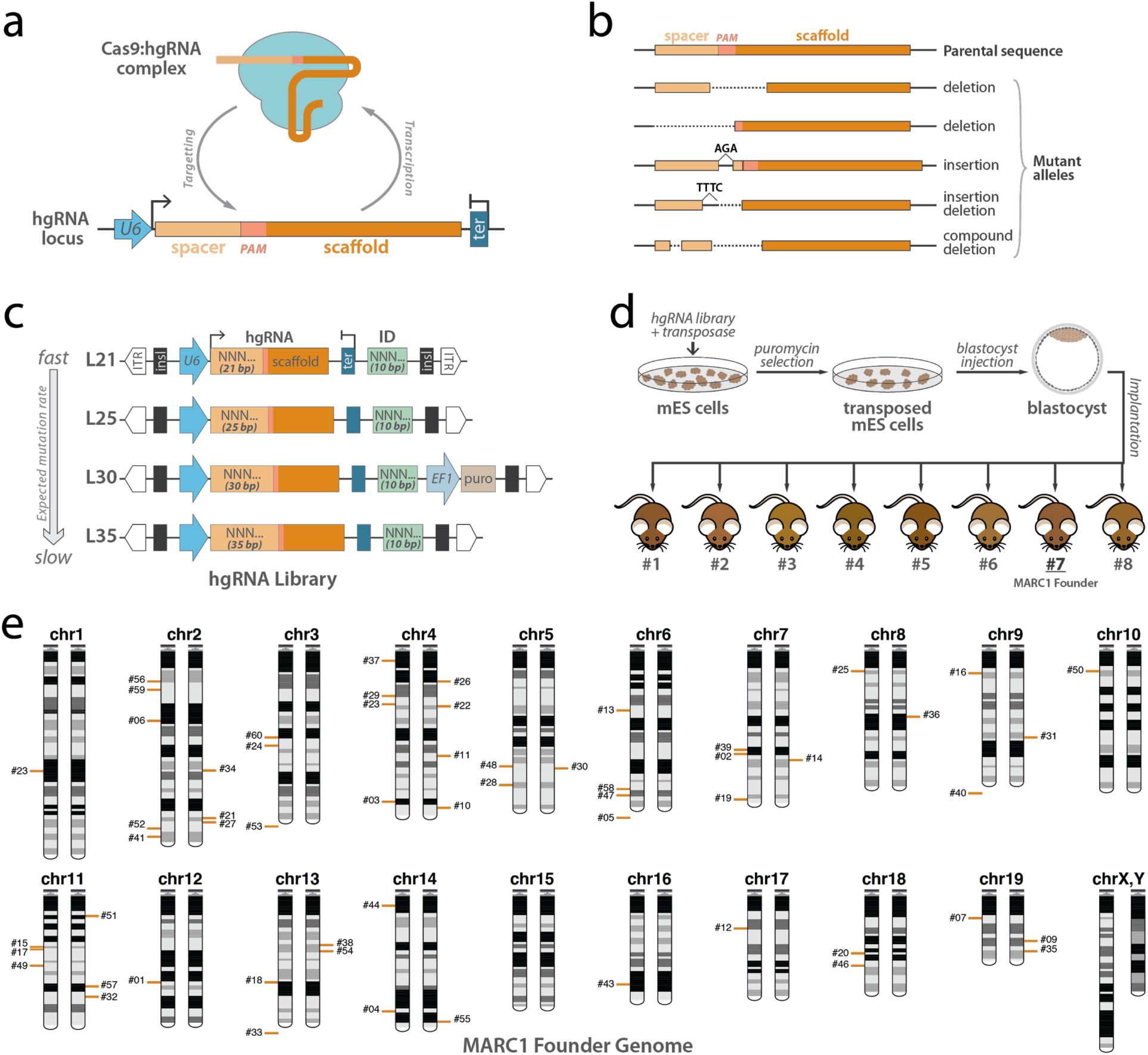
hgRNAs and transgenic strategy to generate mice with multiple hgRNA integrations. **(a)** Homing CRISPR system, in which the Cas9:hgRNA complex cuts the locus encoding the hgRNA itself. As the NHEJ repair system repairs the cut (Lieber & Wilson 2010), it introduces mutations in the hgRNA locus. **(b)** Example of mutations that are created in the hgRNA locus that can effectively act as barcodes. **(c)** Design of PiggyBac hgRNA library for creating transgenic mouse. Four hgRNA sub-libraries with 21, 25, 30, and 35 bases of distance between transcription start site (TSS) and scaffold PAM were constructed and combined. The shortest construct (L21) was expected to have the highest mutation rate with increasing construct length leading to lower mutation rates (Kalhor et al. 2017). The spacer sequence (light orange box) and the identifier sequence (green box) were composed of degenerate bases. Puromycin N-acetyl-transferase gene, which confers resistance to puromycin, was integrated only in the L30 sub-library. **(d)** Blastocyst injection strategy for producing hgRNA mice. The hgRNA library was transposed into mouse ES cells using PiggyBac transposase. Cells with a high number of transpositions were enriched using puromycin selection and injected in E3.5 mouse blastocysts to obtain chimeras. Chimera #7, MARC1 founder, was found to have the highest number of heritable hgRNAs and was chosen as a founder. **(e)** Chromosomal position of all 54 hgRNAs whose genomic position was deciphered in the MARC1 founder (red bars). Bars on left or right copy of the chromosome indicate the hgRNAs that are linked on the same homologous copy. hgRNAs whose exact genomic position is not known but whose chromosome can be determined based on linkage are shown below the chromosome. ITR: PiggyBac Inverted Terminal Repeats; insl: insulator; U6: U6 promoter; ter: U6 terminator; ID: Identifier sequence; EF1: Human elongation factor-1 promoter; puro: puromycin resistance, mES cells: mouse embryonic stem cells.

## Results

### Generation of transgenic founder mouse with multiple hgRNA loci

We created a library of hgRNAs with four different transcript lengths, variable spacer sequences, and 10-base identifiers downstream of the hgRNA scaffold in a PiggyBac transposon backbone (Figure 1c, Materials and Methods). We then transposed this library into mouse embryonic stem (mES) cells under conditions that would result in a high number of transpositions per cell (Figure 1d, Materials and Methods). Transfected mES cells were injected into blastocysts which were then implanted in surrogate females to generate chimeric mice. 23 chimeric mice resulted, of which, eight males were more than 75% transgenic based on their coat color (Figure 1d). Five of the eight showed more than 20 total hgRNA integrations in their genomes and were crossed with wild-type mice. The chimera with the highest average number of hgRNAs transmitted to its progeny, 1763PB7, was chosen for further studies and starting a line. We refer to this mouse as the MARC1 (Mouse for Actively Recording Cells!) founder and its progeny as the MARC1 line. All results from here on focus on the MARC1 founder and its progeny.

### Characterization of hgRNA sequences, positions, and inheritance

By sequencing the hgRNA loci in the MARC1 founder, we identified 60 different hgRNAs (Table 1, Table S1). Each hgRNA has a unique 10-base identifier and a different spacer sequence (Table S1, we will refer to each hgRNA by its ID sequence or number). We also sequenced the regions immediately flanking the transposed elements (Materials and Methods), which allowed us to determined the genomic positions of 54 of the 60 hgRNAs (Table S1, Materials and Methods). We then crossed the MARC1 founder with multiple females and analyzed germline transmission and the inheritance pattern of these hgRNAs in the more than one hundred resulting offspring. All 60 hgRNAs were transmitted through the germline and the offspring carrying them were fertile, had normal litter sizes, and presented no morphological abnormalities. 55 of the 60 showed a Mendelian inheritance pattern, appearing in about 50% of the offspring (Table 1, Table S1). An additional 3 of the 60 were detected in less than 20% of the offspring which we attribute to the low detection rate of L30 hgRNAs due to the performance of the PCR primer used for these and only these three hgRNAs (Materials and Methods). The remaining two were inherited in almost 75% of the offspring, a result best explained by duplication of the hgRNAs to loci more than 50 centimorgans away on the same chromosome or on a different chromosome and confirmed by the genomic location data (Table 1, Table S1, Materials and Methods).

**Table 1.** Summary of all hgRNAs in the MARC1 founder male. For each hgRNA, its ID number, TSS to PAM length (L), observed inheritance probability (Inh%), chromosomal position (“+” indicates transcription in the direction of positive reference strand). See Tables S1, S2, and S3 for more details.

| ID# | L(bp) | Inh% | Position | Class | ID# | L(bp) | Inh% | Position | Class |
| --- | --- | --- | --- | --- | --- | --- | --- | --- | --- |
| 1 | 21 | 49.6 | chr12:84168907:+ | Late | 31 | 35 | 58.4 | chr9:85135810:- | Inactive |
| 2 | 35 | 55.2 | chr7:100292848:+ | Inactive | 32 | 24 | 44 | chr11:98129357:+ | Mid |
| 3 | 35 | 55.2 | chr4:147419769:+ | Inactive | 33 | 21 | 55.2 | chr13 | Late |
| 4 | 21 | 29.6 | chr14:112445289:+ | Inactive | 34 | 35 | 40 | chr2:116850098:+ | Mid |
| 5 | 35 | 41.6 | chr6 | Inactive | 35 | 25 | 52 | chr19:54021355:+ | Late |
| 6 | 21 | 54.4 | chr2:69904041:- | Mid | 36 | 21 | 45.6 | chr8:64294161:- | Mid |
| 7 | 34 | 43.2 | chr19:23587076:- | Late | 37 | 35 | 51.2 | chr4:11578767:+ | Late |
| 8 | 30 | 16.8 |  | Late | 38 | 21 | 38.4 | chr13:48018178:+ | Mid |
| 9 | 35 | 51.2 | chr19:44781921:- | Late | 39 | 25 | 54.4 | chr7:97091435:- | Mid |
| 10 | 25 | 42.4 | chr4:152649298:+ | Late | 40 | 35 | 45.6 | chr9 | Inactive |
| 11 | 25 | 35.2 | chr4:103214352:- | Inactive | 41 | 21 | 38.4 | chr2:181932341:- | Mid |
| 12 | 35 | 48.8 | chr17:32993034:+ | Inactive | 42 | 30 | 15.2 |  | Late |
| 13 | 35 | 42.4 | chr6:59682549:- | Late | 43 | 35 | 48.8 | chr16:87663120:- | Late |
| 14 | 35 | 43.2 | chr7:106645806:- | Inactive | 44 | 21 | 51.2 | chr14:10894128:- | Inactive |
| 15 | 34 | 60.8 | chr11:51639162:- | Late | 45 | 30 | 7.2 |  | Inactive |
| 16 | 35 | 47.2 | chr9:23011278:- | Inactive | 46 | 25 | 47.2 | chr18:68792401:- | Late |
| 17 | 21 | 59.2 | chr11:53401186:+ | Late | 47 | 35 | 48.8 | chr6:140630931:+ | Inactive |
| 18 | 25 | 50.4 | chr13:84138896:+ | Late | 48 | 35 | 34.4 | chr5:113271521:+ | Late |
| 19 | 21 | 39.2 | chr7:145314072:- | Early | 49 | 35 | 58.4 | chr11:69995251:+ | Late |
| 20 | 21 | 49.6 | chr18:57857891:- | Early | 50 | 21 | 44 | chr10:20649492:+ | Mid |
| 21 | 35 | 40 | chr2:163466115:- | Late | 51 | 25 | 36.8 | chr11:21285273:+ | Late |
| 22 | 21 | 42.4 | chr4:55136321:- | Mid | 52 | 21 | 53.6 | chr2:173089614:- | Early |
| 23 | 21 | 80.8 | chr4:53835610:- AND<br>chr1:113070656:+ | Mid | 53 | 35 | 47.2 | chr3 | Inactive |
| 24 | 21 | 51.2 | chr3:93592769:- | Inactive | 54 | 35 | 73.6 | chr13:55598605:+<br>AND chr10:115251722:- | Late |
| 25 | 25 | 48 | chr8:20267522:+<br>OR chr8:20583339:- | Inactive | 55 | 25 | 68.8 | chr14:122192615:+ | Late |
| 26 | 35 | 46.4 | chr4:31943863:- | Inactive | 56 | 21 | 55.2 | chr2:31153437:- | Late |
| 27 | 21 | 39.2 | chr2:167014655:- | Late | 57 | 34 | 44 | chr11:88908785:+ | Inactive |
| 28 | 25 | 40 | chr5:130378620:- | Late | 58 | 21 | 43.2 | chr6:135360506:+ | Early |
| 29 | 35 | 52.8 | chr4:45462426:+ | Late | 59 | 21 | 50.4 | chr2:39102456:+ | Early |
| 30 | 35 | 55.2 | chr5:114521001:- | Inactive | 60 | 25 | 51.2 | chr3:84200519:- | Late |

We also compared the co-inheritance frequencies of these hgRNAs to that expected from Mendelian inheritance of independently segregating loci (Figure S1). We found no mutually-exclusive cosegregating groups of hgRNAs (Figure S1a), indicating that the entire germline in MARC1 founder was derived from only one of the injected stem cells and is thus genetically homogeneous. Considering that every hgRNA detected in the somatic tissue of the MARC1 founder was also inherited to its offspring, these results further suggest that almost all transgenic cells within this chimera were derived from one of the stem cells that were injected into its blastocyst, an observation consistent with previous studies (Markert & Petters 1978; Wang & Jaenisch 2004). The co-inheritance analysis also revealed 13 groups of hgRNAs that deviate from an independent segregation pattern, suggesting that they are linked on a chromosome (Figure S1b). Close examination of this linkage disequilibrium allowed us to determine which linked hgRNAs were on different homologous copies of the same chromosome or were linked on the same copy of a chromosome (Figure S1c). Combined with the genomic location information that was obtained by sequencing, this co-inheritance analysis allowed us to decipher the cytogenetic location of most hgRNAs in the MARC1 founder with a high degree of confidence (Figure 1e).

Based on these results, we concluded that the 60 hgRNAs in the MARC1 line are distributed in almost all chromosomes, are inherited according to a Mendelian pattern, do not interfere with normal animal development and cellular functions in their heterozygote state, and are stable in the absence of Cas9 nuclease.

### Characterization of hgRNA activity

We next studied the activity of these MARC1 hgRNAs upon activation with Cas9. For that, we crossed the MARC1 founder with Rosa26-Cas9 knockin females which constitutively express Cas9 (Platt et al. 2014), effectively delivering Cas9 to the zygote. From this MARC1 x Rosa26-Cas9 cross, 95 offspring at various embryonic stages or the adult stage were obtained (Table 2). While only a single sample was obtained from most offspring, multiple tissues were sampled in some for added statistical confidence (Table 2, Materials and Methods). The hgRNA loci in each sample were then amplified and sequenced as a pool. The ID sequence was used to group the sequencing reads corresponding to each hgRNA and analyze the mutations in its spacer (Figure 2). The results show a wide range of activity among the 60 hgRNAs, with some mutating quickly after Cas9 introduction in almost all cells while some others showed no activity (Figure 2a, Table S2). Five hgRNAs can be considered “early” as they are mutated in at least 80% of the cells in each sample by E3.5 and are mutated in almost all cells by E8.5. Twenty six can be considered “late” as they are mutated in a minority of cells even in the adult stage. Ten more can be considered between “early” and “late” (“mid”) as they continue to mutate throughout embryonic development and are mutated in almost all cells only in later embryonic stages or in the adult animals. The remaining hgRNAs appear to be inactive, at least with this level of Cas9 expression, with a mutation rate below 2% in all samples (Table 1, Table S2).

**Table 2.** A break down of the developmental stage (columns) of all mice used for hgRNA activity analysis and number of samples obtained per mouse (rows). In total, 185 samples from 95 mice were obtained and sequenced.

|  |  | Stage of development |  |  |  |  |  |  |  |  |  | Total mice | Total samples |
| --- | --- | --- | --- | --- | --- | --- | --- | --- | --- | --- | --- | --- | --- |
|  |  | E3.5 | E6.5 | E7.5 | E8.5 | E10.5 | E12.5 | E14.5 | E15.5 | E16.5 | adult |  |  |
| Samples per mouse | one | 11 | 6 | 3 | 3 | 1 | 0 | 0 | 0 | 0 | 38 | 62 | 62 |
|  | two | 0 | 0 | 4 | 6 | 1 | 0 | 0 | 0 | 0 | 0 | 11 | 22 |
|  | three | 0 | 0 | 1 | 0 | 0 | 0 | 0 | 0 | 0 | 0 | 1 | 3 |
|  | four | 0 | 0 | 0 | 0 | 7 | 6 | 0 | 0 | 2 | 0 | 15 | 60 |
|  | five | 0 | 0 | 0 | 0 | 0 | 1 | 0 | 0 | 0 | 0 | 1 | 5 |
|  | six | 0 | 0 | 0 | 0 | 0 | 2 | 0 | 0 | 0 | 0 | 2 | 12 |
|  | seven | 0 | 0 | 0 | 0 | 0 | 3 | 0 | 0 | 0 | 0 | 3 | 21 |
| Total |  | 11 | 6 | 8 | 9 | 9 | 12 | 0 | 0 | 2 | 38 | 95 | 185 |

**Figure 2.**
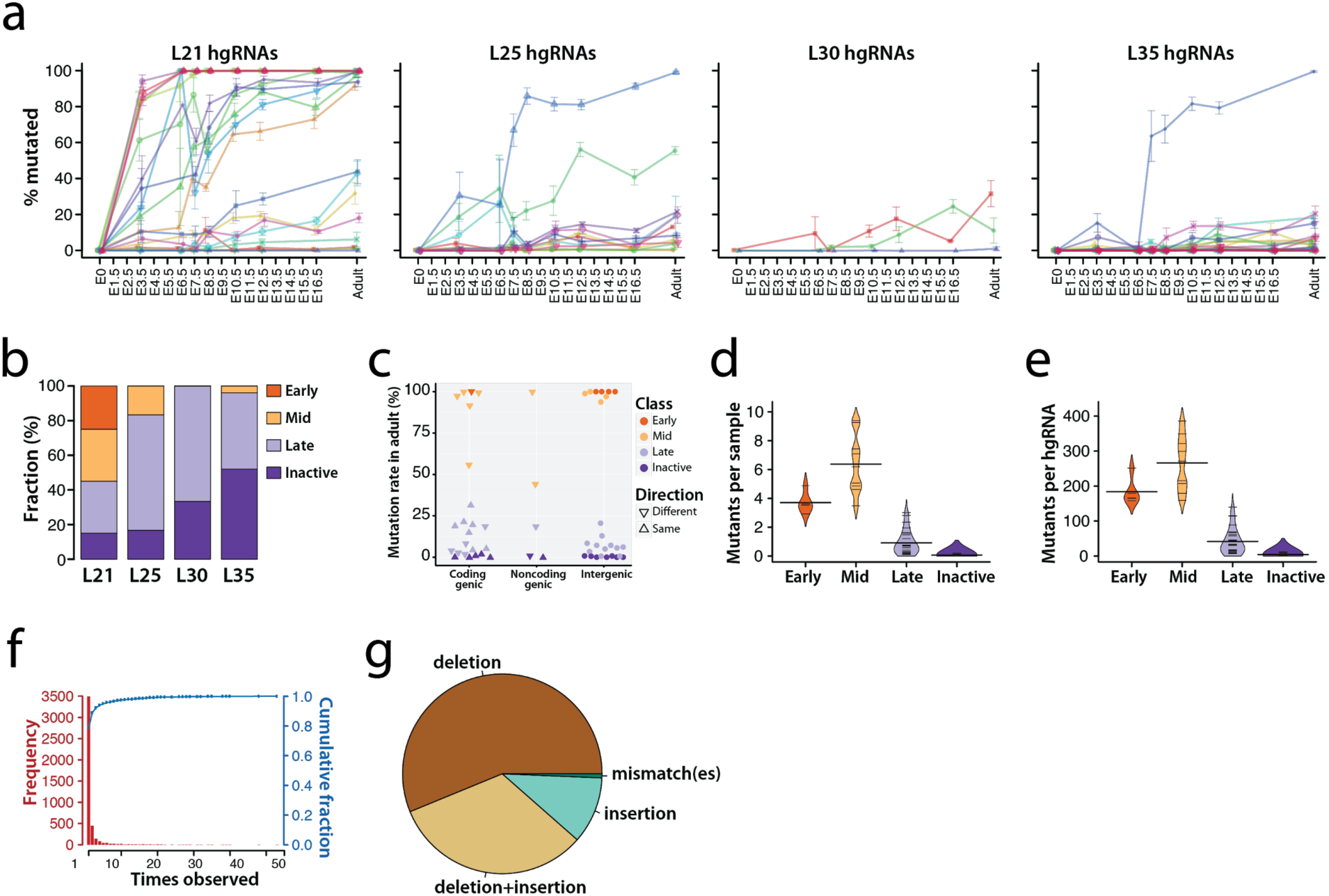
Activity of MARC1 hgRNAs. **(a)** Activity profiles of all 60 hgRNAs in embryonic and adult progenies of MARC1 founder crossed with Cas9-knockin females, broken down by hgRNA length. The fraction of mutant (non-parental) spacer sequences in each hgRNA is measured. Each line connects the observed averaged mutation rates of a single hgRNA. Mean ±SEM is shown, number of samples is different for each value, depending on the number of mouse samples analyzed at the specific time point that had inherited the hgRNA (Table 2). Colors are only meant to distinguish different hgRNAs. See Table S2 for numerical values of the plot. **(b)** Functional categorization of hgRNAs based on their activity profile in panel a, broken down by length. **(c)** Position and transcription direction of hgRNAs with respect to all known coding and non-coding genes, annotated for their functional categorization. See Table S3 for the genes hgRNAs are located in and Figure S2 for breakdown of this plot by hgRNA length. **(d,e)** Distribution of the average number of observed mutant spacers (d) and total number of mutant spacers per mouse (e) for hgRNAs of each category. See Figure S3 for a separate plot for each hgRNA. **(f)** Histogram (red bars) and cumulative fraction (blue connected dots) of the number of mice each mutant spacer was observed in, combined for all hgRNAs. About 90% of all mutant spacers were only observed in one or two mice. See Figure S4 for a separate plot for each hgRNA. **(g)** Types of mutations observed in mutant spacers. See Table S4 for the sequences and alignment of all mutants and Figure S5 for a separate plot for each hgRNA.

Several factors appear to affect hgRNA activity. Transcript length has a clear effect: a far higher fraction of L21 hgRNAs, which have the shortest possible transcript length, are active compared to L25, L30, and L35 hgRNAs which are longer by 4, 9, and 14 bases, respectively (Figure 2a,b). Additionally, only L21 hgRNAs show “early” activity while in longer hgRNAs the inactive proportion seems to grow (Figure 2b). Another factor affecting hgRNA activity levels is their genomic location and interplay with endogenous elements (Table S3). We detected no significant difference between hgRNAs that are in intergenic regions and those within known coding and non-coding genes (Wilcoxon p-value > 0.1, Figure 2c, Figure S2). On the other hand, among hgRNAs that have landed within known coding and non-coding genes, those that transcribe in the same direction as the gene have a lower activity than those that transcribed in the opposite direction (Wilcoxon p-value < 0.01, Figure 2c, Figure S2).

We next analyzed the diversity of the mutations produced by these hgRNAs, which is an important factor in applying them for barcoding and lineage tracing purposes (Kalhor et al. 2017). In a previous study, where hgRNAs were activated in cultured cells and their mutation landscape catalogued without selection, we used entropy as a measure of diversity (Kalhor et al. 2017). In here, where the observed mutations are bottlenecked by the small number of cells early in development and their proportion is skewed by the number of cells that descend from each founder, entropy cannot be straightforwardly measured. In fact, on average, only 4 different mutant spacers were observed for each “early” hgRNA in each sample, and only 6 different mutant spacers were observed for each “mid” hgRNA in each sample (Figure 2d, Figure S3a). However, when combining the results from all animals with a given hgRNA, on average, 200 distinct mutant spacers for each “early” hgRNA and 300 for each “mid” hgRNA were observed (Figure 2e, Figure S3b), suggesting that the hgRNAs can create a large diversity of mutations in mice such that the likelihood of the same mutation occurring independently more than once is small. In fact, for each hgRNA, only a minority of the mutants are observed in more than one animal (Figure 2f, Figure S4), indicating that the diversity space is so large that it cannot be adequately explored with our sampling of 95 animals. The vast majority of this diversity is created by indel mutations (Figure 2g, Figure S5, Table S4) in patterns consistent with those reported for targeting a site with a single gRNA (Shin et al. 2017).

### In vivo barcodes generated by hgRNA

We next analyzed the in vivo barcodes that were generated in hgRNA loci after activation with Cas9 (Figure 3). We focused on eight post E12 embryos from the MARC1 x Rosa26-Cas9 cross for which four different tissues had been sampled (Table 2). The sampled tissues were the placenta, the yolk sac, the head, and the tail. First, we analyzed each “early” and “mid” hgRNA in all the embryos to which it had segregated (Figure 3a,b, Figure S6). For each hgRNA, we identified all mutant alleles observed in these embryos and their relative abundance in each sampled tissue. For each hgRNA, each sample is represented by a mutation frequency vector, or a barcode, comprising a set of all observed mutant alleles and their abundances (Figure 3b, Figure S6). We used the Manhattan distance (L1) to compare the barcodes generated for each hgRNA in all samples in a pairwise fashion (Figure 3c). A scaled distance of 100 between hgRNA barcodes of two samples indicates a completely non-overlapping set of mutant alleles between those samples for the hgRNA locus analyzed. A distance of 0 indicates a complete overlap of mutant alleles with identical relative frequencies. This analysis shows that more than 99% of all barcode pairs have a scaled Manhattan distance of more than 5, indicating unique barcoding of each sample by each hgRNA. Furthermore, barcodes from the tissues of the same embryo were more similar (average distance = 38) whereas those of different embryos were more distinct (average distance = 67) (Figure 3c). These results confirm that hgRNAs undergo random mutagenesis in each embryo to create distinct barcodes in each.

**Figure 3.**
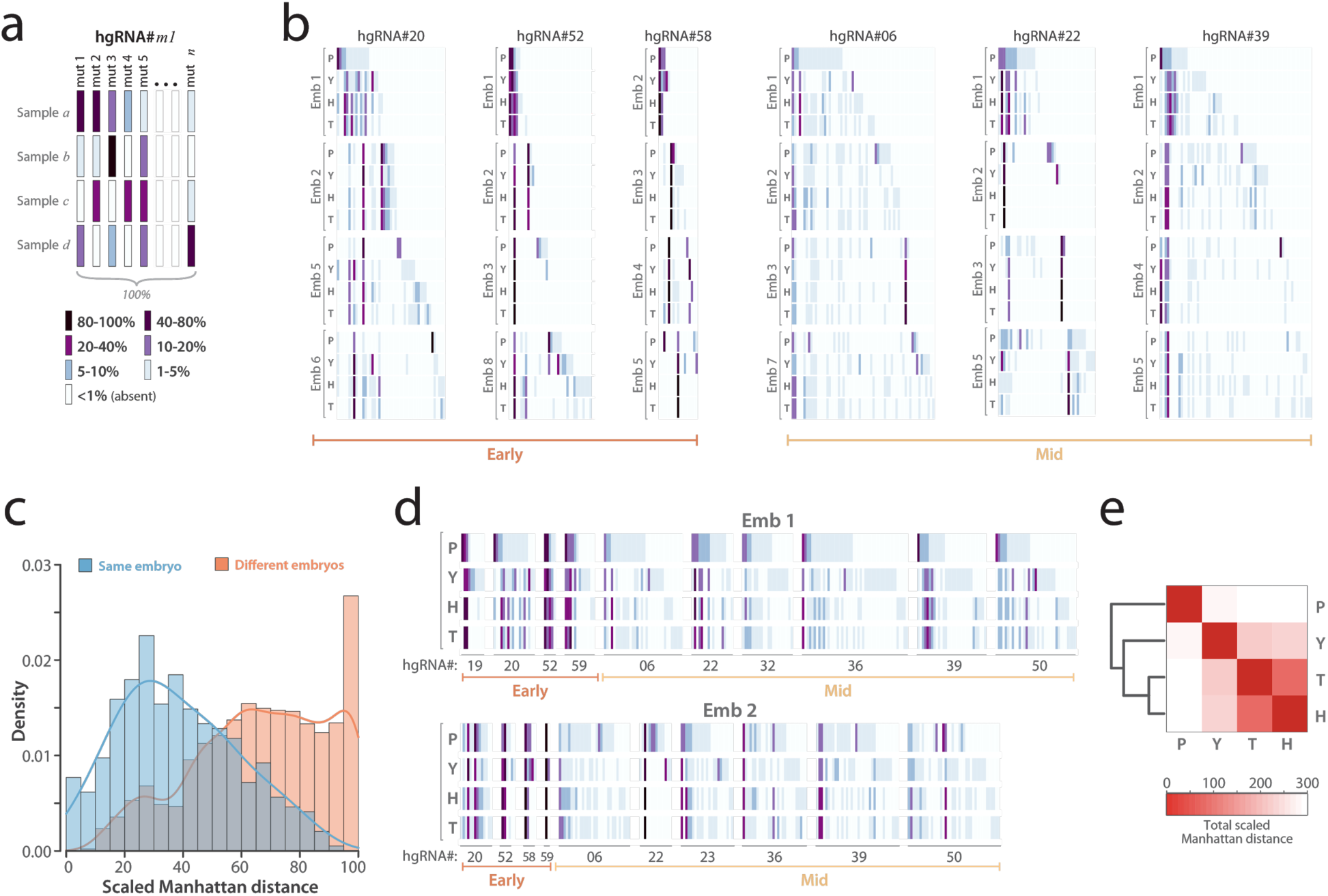
In vivo barcoding in mouse embryos. **(a)** Barcode depiction for each hgRNA in each sample. Each column corresponds to an observed mutant spacer and each row corresponds to a sample. The color of each block represents the observed frequency of the corresponding mutant spacer in the corresponding sample. **(b)** In vivo-generated barcodes of three “early” and three “mid” hgRNAs in eight embryos from a MARC1 x Cas9 cross. Four tissues were sampled from each embryo: the placenta (P), the yolk sac (Y), the head (H), and the tail (T). Embryos 1 and 2 were obtained at E16.5 whereas embryos 3 to 8 were obtained at E12.5 (Table 2). For each hgRNA, the results for a maximum of four embryos is shown. Full barcodes for all hgRNAs in Figure S6. The color code is as annotated in panel a. **(c)** Histogram of the scaled Manhattan distances (L1) between the barcodes of all possible sample pairs for each hgRNA, broken down by sample pairs belonging to the same embryo (blue) and pairs belonging to different embryos (orange). all hgRNA barcodes in all eight embryos **(d)** The complete barcode, comprise of each hgRNA’s barcode, for embryo 1 and embryo 2. **(e)** Heatmap of the average Manhattan distance between the “full” barcodes of placenta, yolk sac, head, and tail samples in all eight embryos. For a separate map for each embryo see Figure S7.

We next asked if the differences between samples of the same embryo reflect their embryonic origin. We created a “full” barcode for each embryo by combining the barcodes generated by each of its hgRNAs (Figure 3d) and compared the Manhattan distance between these barcodes (Figure 3e, Figure S7). These results show higher similarity between the head and tail samples which together are the most different from placenta. Considering that the head and tail samples are derived from the inner cell mass (ICM) whereas the placenta is mostly derived from the trophectoderm (Lu et al. 2001), these results show that hgRNA barcodes of different tissues embody their lineage history. It should be emphasized that an informative lineage analysis may not be carried out on these four samples because they represent overlapping lineages. Specifically, while the head and tail samples are entirely ICM-derived, the yolk sac is comprised of cells from the trophectoderm and the ICM lineages, and the placenta contains trophectoderm-derived cells and a small fraction of cells from the ICM (Lu et al. 2001). Nevertheless, these results suggest that the MARC1 line may be used for tracing lineages in mouse.

## Discussion

Here we describe a mouse strain carrying a scattered array of genomically-integrated hgRNA loci (Figure 1). The array comprises sixty hgRNAs, each with distinct spacer, identification sequence, and genomic location (Table 1). We show that these hgRNAs can be activated to mutate themselves upon introduction of Cas9 nuclease, that different hgRNAs mutate at different rates, and that they produce a large diversity of mutant alleles that is commensurate with the barcoding demands of recording and lineage tracing applications (Figure 2). We further show that barcodes generated in MARC1 x Cas9 cross embryos are compatible with lineage history of various tissues (Figure 3). The extensive benchmarking that has been carried out for each hgRNA in the MARC1 line, together with the ease of crossing it with Cas9 lines, provides an enabling platform for in vivo barcoding in a mammalian model system.

### Mouse line generation and characterization

Our strategy to create the hgRNA transgenic mouse was informed by the required features of an hgRNA mouse as we have previously outlined (Kalhor et al. 2017); namely, the presence of multiple, independent, uniquely identifiable, and genomically integrated hgRNAs that continuously generate diverse barcodes over a stretch of time. Furthermore, we randomly scattered the hgRNA loci in the genome instead of inserting hgRNAs into a single location as a contiguous array, given that the single-location option would easily lead to large deletions or scrambling with multiple adjacent cut sites (Lee et al. 2010; Canver et al. 2014; Byrne et al. 2015), a phenomenon also known as “overwriting.”

To ensure sustained mutagenesis during development, we used hgRNAs of four different lengths in our library, with L21 hgRNAs - the shortest possible length - expected to mutate earliest, and L35 hgRNAs expected to mutate later (Figure 1c). However, surprisingly we found that longer hgRNAs are far less active in mice than expected based on cell culture experiments (Figure 2a,b). In fact, the L35 hgRNAs in mice here seem to be far less active than an L101 hgRNA in cultured HEK/293T cells (Kalhor et al. 2017). Serendipitously, L21 hgRNAs turned out to show a range of activity in mice - from early to late and even inactive - which in turn resulted in sustained mutagenesis through embryonic development (Figure 2a). The reasons for this variation in L21 hgRNA activity are not entirely clear, but positional effects based on hgRNA integration sites appear to play a role (Figure 2c).

### Applications and advantages of MARC1 line

This hgRNA mouse line is a flexible resource that can accommodate disparate demands and varied interests related to in vivo barcoding strategies. An important area will be lineage tracing applications. Using this line, barcoding elements could be activated simply by crossing the MARC1 hgRNA line with any Cas9 line. The Cas9 line will determine when, where, and to what extent in vivo barcoding will take place. For instance, as done here, crossing the MARC1 line with a constitutively active Cas9 initiates in vivo barcoding very early in development and in all cells of mouse, including extraembryonic tissues, without a need for direct injection of barcoding elements to zygotes followed by implantation into surrogate mothers. Almost as easily, one can use other Cas9 lines with higher or lower Cas9 expression, inducible Cas9 expression (Dow et al. 2015; Polstein & Gersbach 2015), or tissue or lineage-specific Cas9 expression.

The other area of focus will be recording cellular signals over time (Perli et al. 2016; Sheth et al. 2017; Marblestone, Zamft, et al. 2013; Shipman et al. 2017; Punthambaker & Hume 2014). Such applications will depend on coupling Cas9 activity with the signal(s) of interest. Yet another application would be to uniquely barcode each cell in a tissue or an organism for identification purposes. In this case, barcodes can be independently generated in each cell by postmitotic activation or delivery of Cas9. Such barcoding applications, particularly for connectome mapping in the brain, are a subject of intense interest (Cai et al. 2013; Marblestone, Daugharthy, et al. 2013; Marblestone et al. 2014; Kebschull et al. 2016; Peikon et al. 2017).

For all these strategies, any upgrades or modifications to the Streptococcus pyogenes Cas9 system, such as more specific versions (Slaymaker et al. 2016; Guilinger et al. 2014; Davis et al. 2015; Kleinstiver et al. 2016) or base-editing Cas9s (Komor et al. 2016; Gaudelli et al. 2017) can be readily combined with this mouse line without a requirement to generate new hgRNA lines.

### MARC1 line maintenance and sharing

Maintenance of this hgRNA mouse line in the future will be similar to other transgenic lines that are maintained in a heterozygote state as breeding all 60 hgRNAs, or even the 41 active ones, into complete homozygosity will take many generations (Haldane & Waddington 1931; Samal & Martin 2017). Furthermore, even though none of the genic hgRNAs were located in an exon (Table S3) or otherwise appear to disrupt normal function of a gene, it is possible that some of these hgRNA loci can result in phenotypic defects in a homozygous state. With these considerations in mind, to start the MARC1 line, we first crossed its founder with multiple wild-type females to obtain a first generation (F1). To obtain the following generation (F2), we genotyped all F1 progeny, and arranged crosses between multiple F1 pairs to maximized number of hgRNAs per F2 progeny, with priority being given to early, mid, and late hgRNAs in that order, followed by homozygosity in those hgRNAs with the same order. The same strategy was applied to F2 to obtain F3, and so on to continue the line. We have established protocols and analysis tools that allow one person to genotype more than one hundred progeny in each generation and determine optimal pairings for creating the next generation in three days, with less than 12 hours of hands-on work, and about $500 in total reagent costs. As we continue breeding the MARC1 line, mice with a determined subset of the hgRNAs are available to the scientific community. As a result, other groups can readily establish and maintain a colony.

## Materials and Methods

### Vectors

Two vectors were constructed based on the PB-CMV-MCS-EF1α-Puro PiggyBac Vector (System Biosciences PB510B-1) and deposited on Addgene. In one vector, PB-U6insert (Addgene plasmid #104536), a U6 promoter with a Multiple Cloning Site (MCS) to its downstream is flanked by core insulators and inverted terminal repeat elements. In the other vector, PB-U6insert-EF1puro (Addgene plasmid #104537), in addition to the U6 promoter and its MCS, there is an EF1 promoter driving the expression of Puromycin N-acetyl-transferase gene downstream. Super PiggyBac Transposase vector was obtained from System Biosciences.

### hgRNA library design and construction

Four oligonucleotides encoding four hgRNAs of different lengths and randomized sequence, with degenerate bases in the spacer and identifier, flanked by AgeI and EcoRI restriction sites and primer binding sites were designed (L21, L25, L30, and L35 oligonucleotides in Supplementary notes). In the degenerate spacer region, a partially degenerate base specifying A, C, and G and excluding T (V) was used every fourth base to eliminate the possibility of long poly thymidines stretches which would prematurely terminate transcription by RNA Polymerase III. This arrangement of degenerate bases results in 16,760,438,784 possible different spacer sequences (452,984,832 for L21, 5,435,817,984 for each of L25, L30, and L35) thus making it highly improbable that a transfected cell would receive hgRNAs with identical or even similar spacer sequences that would result in cross reactivity between them. The identifier sequence was comprised of ten fully degenerate bases resulting in 1,048,576 possible sequences. The identifier was positioned downstream of the hgRNA scaffold to facilitate identification of each hgRNA even if all of its spacer was deleted during activation with Cas9.

Each designed oligonucleotide was synthesized using a commercially available service (Integrated DNA Technologies, Inc.) to obtain a diverse mixture of oligonucleotides. Each mixture was amplified in a PCR reaction with LibPrimF and LibPrimR primers (Supplementary notes) and digested with AgeI and EcoRI. Digested products of L21, L25, and L35 were ligated into the AgeI-EcoRI-digested PB-U6insert vector, whereas digested product of L30 was ligated into the AgeI-EcoRI-digested PB-U6insert-EF1puro vector, using Quick Ligation Kit (NEB). Each ligated product was transformed in ElectroMAX Stbl4 competent cells (Thermo Fisher Scientific). Roughly 100,000 colonies were obtained per transformation compared to less than 1,000 colonies for no-insert controls. All colonies belonging to the same ligation were combined and subjected to plasmid extraction. The L21, L25, L30-puro, and L35 plasmid libraries were mixed at a 6:6:1:6 respective molar ratio to obtain the final hgRNA library.

### Mouse embryonic stem (mES) cells

V6.5 mES cells, established from the inner cell mass (ICM) of an Agouti male E3.5 blastocyst from a C57BL/6 X 129/Sv cross (Eggan et al. 2001), were cultured on a DR4 Mouse Embryonic Fibroblasts (Amsbio, LLC) feeder layer in mES culture medium (DMEM supplemented with 15% ES Cell Qualified Fetal Bovine Serum, 1X nucleosides, 1X Penicillin-Streptomycin, 1X non-essential amino acids, 2 mM L-glutamine, 1X 2-mercaptoethanol, and 1000 units/mL leukemia inhibitory factor, all Millipore-Sigma Embryomax reagents used per manufacturer’s instructions). When necessary, selection was carried out in presence of 1 μg/mL puromycin.

### mES transfection

Transfections were carried out by electroporation using the Lonza P3 Primary Cell 4D X Kit and 4D Nucleofector per manufacturer’s instructions. To obtain a high number of integrated hgRNAs per cell, we used a very high ratio of transposon to transposases during transfection into the mES cells, and we integrated the selectable puromycin marker in only about 5% of all transposons. As such, after puromycin selection, the mES cell population was enriched for a high number of transpositions. Specifically, 4 million dissociated mES cell, 7 μg of the final hgRNA PiggyBac vector library, and 0.7 μg of Super PiggyBac Transposase vector were resuspended in 100 μl of P3 nucleofection solution, transferred to a Nucleocuvette, transfected with program CG-104, and transferred to a well with fresh mES culture medium and feeder cells. Puromycin selection was applied starting 48 hours after transfection and continued for 5 days, at which point cells were cryopreserved. Extensive cell death was observed after puromycin selection. Culture media was renewed the day after nucleofection and each day after puromycin selection. During this time, cells were passaged when ∼70% confluent.

### Blastocyst injection and implantation

Blastocyst injection and implantation into pseudopregnant mothers were carried out by the Genome Modification Facility at Harvard University, using standard procedures (Behringer et al. 2013). Briefly, puromycin-selected stem cells were injected into 30 blastocysts of a C57BL/6J background and implanted into 3 pseudopregnant recipients. This process resulted in 23 viable chimeric mice. Based on their coat colors (agouti donor injected in black recipient), four of the chimera were more than 95% transgenic males, four were 80-90% transgenic males, eight others were 10-70% transgenic males, and the rest were between 5 to 60% transgenic females.

### Animals

All animal procedures were approved by the Harvard University Institutional Animal Care and Use Committee (IACUC). C57BL/6J strain (Stock 000664) and Rosa26-Cas9 knockin strain (Stock 024858) (Platt et al. 2014) were obtained from The Jackson Laboratory. For embryonic samples, an hgRNA male was crossed with Rosa26-Cas9 knockin females. Pregnant females were dissected at the desired embryonic timepoints, designating noon of the day of vaginal plug detection as E0.5.

### Sampling and dissections

For postnatal and adult animals, samples were collected from tail tips or ear clips. Dissection of the embryos was carried out as described (Behringer et al. 2013), and either the entire embryo or the desired regions were sampled for DNA extraction (see Table 2).

### DNA extraction

For genotyping purposes, genomic DNA was extracted from animal tail or ear clips using Qiagen DNeasy Blood and Tissue Kit. For determining hgRNA mutations in Cas9 activated samples, Qiagen DNeasy Blood and Tissue Kit was used when an adequately large tissue sample could be obtained. In cases where only a small number of cells were collected, such as the E3.5 embryos, hgRNA loci were pre-amplified directly from the cells using a protocol derived. For that, each sample was transferred to a 0.2mL tube containing 9 μL of water, 1μL of 10X Cell Lysis and Fragmentation Buffer (Sigma-Aldrich L1043), and 40 mU of Proteinase K (Sigma-Aldrich P4850) and mixed thoroughly. The tube was then incubated at 50 °C for 60 minutes followed by 96 °Cfor 2 minutes. Next, 2 μL of 5 mM Phenylmethanesulfonyl fluoride in 5% ethanol solution (PMSF, Sigma-Aldrich 93482) was added and incubated at room temperature for 15 minutes to inactivate the protease. Finally, the entire 12 μL mix was amplified in a 60 μL PCR with 0.2 μM each of trc-PBLib-F, trc-PBLib-R, and trc-PBLib30R primer for 20 cycles. For these samples, the product of this PCR amplification would serve in place of genomic DNA for later amplification steps.

### hgRNA amplification and sequencing

The hgRNA loci were amplified from genomic DNA templates (or pre-amplified hgRNAs) and prepared for Illumina sequencing using two consecutive PCR amplifications. In the first amplification, between 0.25 and 25ng of extracted genomic DNA was amplified in a 10 μL real-time PCR reaction with 0.2 μM each of the SBS3-PBLib-F and SBS9-PBLib-R, and 0.05 μM SBS9-PBLib30-R primers (Supplementary notes). This amplification was carried out in a real-time setting and stopped in mid-exponential phase, typically after about 25-28 cycles. To add the complete Illumina sequencing adaptors, this first PCR product was diluted and used as the template for a second PCR with NEBNext Dual Indexing Primers, with each sample receiving a different index. In some cases, an in-house set of indexing primers was used to expand the indexing space. Once again, PCR was carried out in a real-time setting and stopped in mid-exponential phase, which was after about 15 cycles in most cases. Samples were then combined and sequenced using Illumina MiSeq with reagent kit v2 or v3. Sequencing was done in both directions, with 200 bases for the forward/first read and 50 bases for the reverse/second read. This amplification and sequencing strategy targets a 220-240 bp amplicon in the hgRNA loci that covers the entire hgRNA, its identifier, 74 bases of the U6 promoter immediately upstream of the spacer, and just over 20 bases immediately downstream of the identifier (Supplementary notes). The first/forward sequencing read reliably covers the hgRNA spacer while the second/reverse read reliably covers the identifier. Total number of samples per run were adjusted to obtain at least 10,000 reads per sample.

### hgRNA sequence analysis

Analysis was carried out based on our previous work (Kalhor et al. 2017) but with modifications. For each sample, the spacer and identifier sequences were extracted from each paired-end read. To obtain the spacer sequence, the 73 base signature “atggactatcatatgcttaccgtaacttgaaagtatttcgatttcttggctttatatatcttgtggaaaggac,” which covers the U6 promoter region immediately upstream of the spacer (Supplementary Notes), was aligned to the forward read and the 50 bases to its immediate downstream were extracted. To obtain the identifier, the 12 base signature “gggcccgaattc,” which covers the region immediately downstream of the identifier (Supplementary Notes), was aligned to the reverse read and the 10 bases to its immediate upstream were extracted. Alignments were carried out using BLAT (Kent 2002) on an in-house server. An 80% identity cutoff was applied: paired-end reads matching less than 80% of either signature sequence were discarded. Combining all the pass-filter reads for each sample, the total observed counts of each unique spacer-identifier pair was tabulated. To adjust for sequencing and amplification error, which inflate the number of unique spacer-identifier pairs by introducing random point mutations, we employed a program that reduces such errors (Beltman et al. 2016). Notably, we modified the code to use the Hamming distance instead of its default Levenshtein distance when comparing spacers because Hamming distance preserves deletions, which are present in our datasets when Cas9 is activated. The final output of this hgRNA sequence analysis is a list of unique spacer-identifier pairs observed in each sample. The list also includes the number of times each pair was observed in the sequencing results. For non-Cas9-activated samples, this list represents their genotype - or hgRNAs that are present in that sample. For Cas9-activated samples, it also shows the mutations that have accumulated in each spacer as multiple spacers for a single identifier.

Importantly, despite filtering steps described above, the lists tend to contain various erroneous spacer-identifier pairs that are observed at far lower frequencies than the true spacer-identifier pairs, typically 100 to 1,000 times less frequent. These erroneous pairs persist for several reasons. First, not all sequencing errors can be computationally adjusted as described by Beltman and colleagues. Second, template switching and recombination during PCR amplification (Potapov & Ong 2017), which happen at very low rates, can result in spurious association of an identifier with the wrong spacer sequence. Third, index switching during Illumina library preparation and sequencing of multiplexed libraries can create virtual cross-contamination between different samples in a pool (Schnell et al. 2015; Sinha et al. 2017), thus resulting in a low-frequency appearance of spacer-identifier pairs from the rest of the pool in each sample. We focused our analyses on spacer-identifier pairs whose presence could be determined with a high confidence level. Specifically, from all the identifiers observed in each sample, we removed any identifiers whose total abundance, irrespective of spacer, was less than 0.1% of all identifiers with a higher abundance. Furthermore, for each identifier, we removed spacers whose frequency was less than 0.01% of all spacers observed for that identifier. It should be noted that, depending on the quality of sequencing runs and the total number of identifiers expected for a sample, slightly higher or lower cutoff values may have been applied. However, in all cases, the appropriate cutoffs could be determined with confidence due to the large drop in frequency of apparently erroneous pairs compared to the more frequent ones. This final filtering step resulted in a list of high-confidence unique spacer-identifier pairs observed in each sample and their respective abundances.

### Genomic location of hgRNAs

It was determined that three 4-cutter restriction enzymes CviQI, HpyCH4III, and TaqQI, do not have any putative restriction sites in the 564 bp region between the identifier and 3’-ITR of L21, L25, and L35 hgRNAs. Therefore, digesting MARC1 founder genomic DNA with any of these enzymes will produce DNA fragments which span the identifier and the insertion point of the PiggyBac insert on the 3’-ITR side. These fragments can then be sequenced to identify insertion points in genomic DNA. To implement this strategy, purified MARC1 founder DNA was digested separately with each of CviQI, HpyCH4III, and Taq^α^I restriction enzymes (all from NEB) using the recommended conditions. Double-stranded DNA adaptors CviQI, HpyCH4III, and TaqQI, compatible with CviQI, HpyCH4III, and Taq^α^I-generated overhangs respectively, were constituted by annealing the appropriate single-stranded DNA oligos (Supplementary Notes) in a Tris-EDTA (TE) buffer containing 50mM NaCl. 100ng of each digested genomic DNA sample was then ligated to its corresponding adaptor in 20 μL of 1X Enzymatics Rapid Ligation with 0.6 μM of the adaptor and 600 units of T4 DNA Ligase at room temperature for 20 minutes. Each ligated product was then purified and amplified in a PCR reaction with SBS3-scaffold-F and SBS9-Adaptor-R primers (Supplementary Notes) for 22 cycles. The PCR products between 600 and 1000 bp were then extracted from a 2% agarose gel. To add the complete Illumina sequencing adaptors, this gel-extracted PCR product was used as the template for a second PCR with NEBNext Dual Indexing Primers, with each sample receiving a different index. Samples were then combined and sequenced using Illumina MiSeq with reagent kit v2 with 150 bases for the first read and 150 bases for the second read. The first reads were then used to identify the hgRNA and the second reads were aligned to mouse genome (GRCm38/mm10 assembly) using Bowtie 2 (Langmead & Salzberg 2012) to identify the genomic location of the hgRNAs. Of the 57 L21, L25, and L35 hgRNAs, 40 were located with a “high” confidence (Table S1), meaning that same location was identified for them in at least two of the CviQI, HpyCH4III, and Taq^α^I-generated libraries with a high number of reads. Another 12 were located with “medium” confidence, meaning that their location was identified in one of the three libraries with a high number of reads or two of the three with a low number of reads. Another 2 were located with “low” confidence, meaning their locations were only identified in one of the three libraries with a low number of reads. The remaining 3 could not be located in any of the three libraries due to the repetitive nature of sequences surrounding them or a lack of adjacent CviQI, HpyCH4III, or Taq^α^I site downstream of their insertion point. This location identification scheme is not compatible with the L30 inserts, whose locations were also not identified (Table S1). To find the potential endogenous elements in which our hgRNAs might have landed, we queried the UCSC Genes track of the UCSC Table Browser with the obtained genomic locations (Table S1). Using the knownGene table, we labeled the hgRNAs that are not within a gene as “intergenic”. For the remaining hgRNAs which reside in genes, we labeled those that are in protein coding genes as “coding genic” and those in non-coding genes as “non-coding genic”. No barcode was found to be within an exon; thus all of the “coding genic” hgRNAs are within introns (Table S3).

### Analysis of inheritance and linkage

All F1 progeny of the MARC1 founder, be it in a cross with wt or Cas9 females (Table 2), were considered. For mice with multiple samples available, only one randomly-chosen sample was included. Inheritance probability for each hgRNA *i* (*P*_*i*_) was determined as the fraction of progeny in which the hgRNA was detected. Coinheritance enrichment of hgRNAs *i* and *j* was defined as:

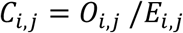

where *O*_*i,j*,(_ is the observed fraction of the times the hgRNAs *i* and *j* were inherited to the same progeny and *E*_*i,j*_ is the expected fraction of progeny to inherit both hgRNAs assuming their independent segregation and is calculated as:

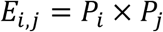

To allow better visualization and analysis of linkage without concern for positioning of hgRNAs on the same or different copies of homologous chromosomes, Linkage index of hgRNAs *i* and *j* was defined as:

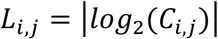

## Acknowledgements

The authors would like to acknowledge Drs. Rudolf Jaenisch and Yonatan Stelzer for helpful discussions and for generously sharing reagents, Dr. John Aach and Leo Meijia for critical reading of the manuscript and helpful comments, Andyna Vernet for assistance with animal husbandry, Dr. Lin Wu and the Genome Modification Facility for blastocyst injections as well as helpful discussions, Dr. Scott Kennedy, Brandon Fields, and Angela Reslow for generously sharing equipment and providing technical advice, and Alex Ng and Nick Conway as well as Drs. John Young, Richie Kohman, Babak Khalaj, Matthew Warman, and Noah Davidsohn for helpful discussions. This work has been supported by funding from NIH grants MH103910 and HG005550 (G.M.C.) and the Intelligence Advanced Research Projects Activity (IARPA) via Department of Interior/Interior Business Center (DoI/IBC) contract number D16PC00008 (G.M.C.).

## Competing Financial Interests

The authors declare no competing financial interest. GMC’s competing financial interests can be found at v.ht/PHNc.

